# Strain-specific differences in the response to egg-derived versus recombinant protein influenza vaccines

**DOI:** 10.64898/2026.02.23.707528

**Authors:** Andrea N. Loes, Rosario Araceli L. Tarabi, Shuk Hang Li, Reilly K. Atkinson, John Huddleston, Caroline Kikawa, Tachianna Griffiths, Elizabeth M. Drapeau, Sook-San Wong, Samuel M. S. Cheng, Nancy H.L. Leung, Sarah Cobey, Benjamin J. Cowling, Trevor Bedford, Scott E. Hensley, Jesse D. Bloom

**Affiliations:** Howard Hughes Medical Institute, Seattle, WA; Division of Basic Sciences and Computational Biology Program, Fred Hutch Cancer Center, Seattle, WA; Department of Microbiology, Perelman School of Medicine, University of Pennsylvania, Philadelphia, PA; Vaccine and Infectious Disease Division, Fred Hutch Cancer Center, Seattle, WA; Department of Genome Sciences, University of Washington, Seattle, WA; Medical Scientist Training Program, University of Washington, Seattle, WA; World Health Organization Collaborating Centre for Infectious Disease Epidemiology and Control, School of Public Health, The University of Hong Kong, Hong Kong, SAR, China; Department of Ecology and Evolution, University of Chicago, Chicago, Illinois, USA

## Abstract

The 2023/2024 influenza vaccine included an updated H1N1 component designed to better match a new clade of H1N1 that had multiple mutations in antigenic epitopes of hemagglutinin. Despite this update, the vaccine trended towards being less effective against the vaccine-matched H1N1 clade than the parental H1N1 clade lacking the new antigenic mutations. Here we measure neutralization titers of serum antibodies from individuals who had received either a recombinant protein or an egg-derived vaccine against a set of viruses with hemagglutinins from 58 H1N1 strains representative of the diversity during the 2023/2024 season. We find that egg-derived vaccine recipients, but not recombinant protein vaccine recipients, had a relatively lower boost in neutralizing titers to the new clade that the updated vaccine was designed to target. We suggest that the difference in the extent that the egg-derived versus recombinant protein vaccines boosted neutralizing titers to the new H1N1 clade is because the seed strain for the egg-derived vaccine strain had acquired a reversion of a key antigenic mutation (K142R) present in that clade. Our results show how egg-derived versus recombinant protein vaccines can elicit different relative titer boosts against different subsets of viral strains, a phenomenon that could impact vaccine effectiveness.

**Importance:** Influenza vaccines can be produced from virus grown in eggs, or grown in cells or made with recombinant protein. Egg-derived influenza vaccines often contain egg-adaptive mutations in the viral antigen hemagglutinin (HA) which can impact the antigenicity or immunogenicity of the HA. In this study, we compare neutralization titers from egg-derived and recombinant protein vaccine recipients against recently circulating influenza A(H1N1) strains. We find that the egg-derived vaccine induces less of a boost in titers than the recombinant protein vaccine to the new clade of viral strains that the vaccine was designed to target.

## Introduction

Twice per year, recommendations are made about which viral strains to include in updated influenza vaccines for the Northern or Southern Hemispheres^1^. These vaccine strains are selected to be antigenically similar to viruses that are predicted to predominate in the following season^2,3^. The majority of influenza vaccine antigens are produced through propagation of viruses in eggs, although some vaccine antigens are produced from viruses grown in cells or recombinantly expressed viral hemagglutinin^4^. As there are limitations in which viral strains grow well in eggs that typically do not apply to vaccines produced in cells, independent strains for each subtype are recommended for the cell-derived and recombinant protein vaccines versus the egg-derived vaccines. Generally, the seed strain for the egg-derived vaccine is a reassortant virus that has been passaged in eggs to improve its growth^4^. Unfortunately, influenza viruses selected for the egg-derived vaccine frequently contain mutations within the hemagglutinin (HA) protein that may improve viral growth in eggs, but also alter antigenicity or immunogenicity^4^. These mutations can alter the effectiveness of the resulting vaccine, particularly if these mutations are in epitopes where recent antigenic changes have occurred within circulating strains^5–10^.

Here we used a high-throughput sequencing-based neutralization assay^11–13^ to examine the change in serum neutralizing antibody titers against recently circulating human H1N1 strains among individuals that received an egg-derived or recombinant protein 2023/2024 vaccine. We find substantial heterogeneity in the strain-specific neutralizing antibody response among individuals within both groups of vaccinees, but even within this heterogeneity it is apparent that the egg-derived vaccine elicited a poorer neutralizing antibody titer boost than the recombinant protein vaccine against the new H1N1 clade that the vaccine was designed to target.

## Results

### Update of the 2023/2024 H1N1 component of the influenza vaccine

The H1N1 strains chosen for inclusion in the 2023/2024 season vaccines were selected from an emerging clade with many antigenic mutations compared to the prior year’s vaccine (Figure 1A)^14^. This new clade, termed 5a.2a.1, had several mutations within known antigenic epitopes (Figure 1B)^15^. This 5a.2a.1 clade, which now accounts for nearly all of the H1N1 influenza found in humans globally (Figure 1C), has circulated at high levels in the United States since its emergence in 2022 (Figure 1D). During the 2023/2024 influenza season, 5a.2a.1 accounted for approximately 75% of the sequenced H1N1 influenza in the United States, and 41% of the sequenced H1N1 influenza globally (as assessed by sequences uploaded to GISAID and consistent with CDC reports^16,17^).

**Figure 1.**
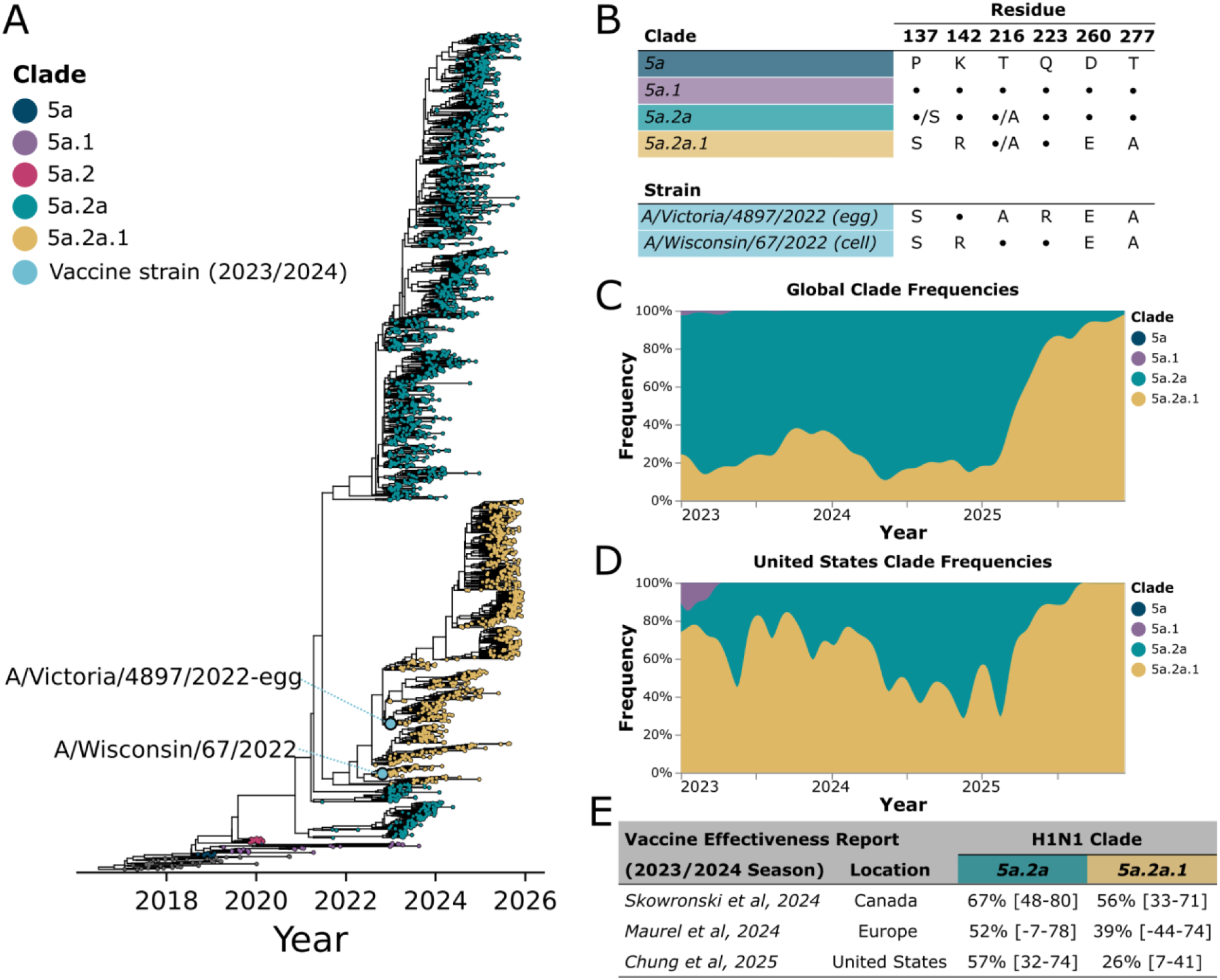
H1N1 vaccine strains for 2023/2024 season were selected from a newly emerging subclade designated 5a.2a.1. A) Phylogeny of HAs from human H1N1 strains circulating from 2022-2025, adapted from the Nextstrain (https://nextstrain.org/seasonal-flu/h1n1pdm/ha/3y@2026-01-02) phylogeny^41,42^. Tips are colored according to clade designation and vaccine strains that were selected for the 2023/2024 season are labelled and indicated with blue stars. B) Variable residues between predominant clades and vaccine strains, showing both the vaccine strains used for egg-derived vaccines and the cell- and recombinant-protein vaccines. A • indicates the residue has the same identity as in the 5a clade. Some variability at these residues also exists within subclades; if a given mutation was present at greater that 20% among sequences in that clade in last three years, this is indicated as •/X. Residue numbers are in the H1 numbering scheme. C), D) H1N1 normalized monthly clade frequencies globally and in the United States from 2023-2025. E) H1N1 Clade specific vaccine effectiveness for the 2023/2024 season, with 95% confidence intervals from recent studies^18–20^.

Despite the choice of 5a.2a.1 clade strains as the H1N1 component of the vaccine, vaccine effectiveness for H1N1 tended to be lower against vaccine-matched 5a.2a.1 clade strains than against other strains that circulated in the 2023/2024 season (Figure 1E)^18–20^. This trend of clade-specific effects on vaccine effectiveness, though not statistically significant, was observed in independent vaccine effectiveness studies from multiple geographic locations^18–20^.

The high-growth reassortant (IVR-238) of the egg-derived vaccine strain contained a reversion at one of the key antigenic mutations that was present in the 5a.2a.1 clade, K142R (H1 numbering, K145R in H3 numbering), so that the vaccine strain contains a lysine at this position, while sequences within this clade contain arginine^19^. In addition, the egg-derived vaccine strain contained a common egg-adaptive mutation Q223R (H1 numbering, Q226R in H3 numbering) (Figure 1B)^21–24^. Both of these mutations are within or adjacent to the same antigenic epitope (Ca2), and could have reduced the antigenic match between the egg-derived vaccine and circulating 5a.2a.1 clade strains. The vaccine strain used for cell- and recombinant-protein vaccines did not contain these mutations (Figure 1B), making it more similar to the circulating 5a.2a.1 viruses in this antigenic epitope. Most of the individuals monitored in surveillance network-based vaccine effectiveness studies likely received egg-derived vaccines (which are more widely used in general), and the available 2023/2024 vaccine effectiveness studies do not stratify effectiveness by vaccine type^18–20^. Therefore, whether egg-adaptive mutations played a role in reducing H1N1 vaccine effectiveness in the 2023/2024 season is unclear.

### The egg-derived but not the recombinant protein vaccine elicits relatively lower titers to strains within clade from which vaccine was selected

To determine if there was a difference in neutralization of circulating H1N1 strains in individuals receiving egg-derived or recombinant protein vaccines, we assessed the specificity of the neutralizing antibody response in individuals vaccinated in the 2023/2024 season by measuring neutralization titers pre- and post-vaccination against a set of 58 H1N1 strains representative of the viruses circulating in that season (Supplemental Figure 1A). We made these measurements using a previously described sequencing-based neutralization assay^11–13^ that makes it possible to simultaneously measure neutralizing titers against many strains at once (Supplemental Figure 1B).

We applied this sequencing-based neutralization assay to sera collected pre- and post-vaccination from adults who received either the egg-derived vaccine FluLaval, or the recombinant protein vaccine Flublok (Table 1). FluLaval is a split-virion, quadrivalent vaccine prepared from viruses grown in eggs, then inactivated, purified by centrifugation, and finally disrupted with detergent^25^. Flublok is a quadrivalent vaccine containing recombinant HA proteins produced in insect cells and purified by column chromatography and has three-times the concentration of HA antigen as the FluLaval vaccine^26^. The cohorts consisted of individuals of similar median age and history of vaccination but from different ethnicities and geographic locations: the egg-derived vaccine cohort consisted of individuals in the USA (Pennsylvania), while the recombinant protein vaccine cohort consisted of individuals from Hong Kong (Table 1). In addition, there may be other differences between these groups that could conceivably represent unknown confounders for our study.

**Table 1.** Overview of vaccinated adult cohorts. Summary of sera from vaccinated adult cohorts that received either the egg-derived or recombinant protein Northern hemisphere influenza vaccine 2023/2024.

|  | egg-derived vaccine recipients | recombinant protein vaccine recipients |
| --- | --- | --- |
| *Number of individuals* | 38 | 37 |
| *Vaccine received* | Northern Hemisphere 2023/2024, egg-derived vaccine (FluLaval) | Northern Hemisphere 2023/2024, Recombinant vaccine (Flublok) |
| *Median (interquartile range, range)* | 1985 (20, 1949-2001) | 1986 (14, 1976-2000) |
| *Location* | Philadelphia, PA (USA) | Hong Kong |
| *Samples collection timing* | 0 and ~28 days post-vaccination | 0 and ~28 days post-vaccination |
| *Prior vaccinations in 2021/2022 and 2022/2023 (Number of participants)* | None (6), 2021/2022 only (3), 2022/2023 only (2), 2021/2022 and 2022/2023 (27) | None (6), 2021/2022 only (0), 2022/2023 only (8), 2021/2022 and 2022/2023 (23) |

Vaccination tended to elicit increases in neutralizing titers against all strains at 28-30 days post-vaccination for both the egg-derived and recombinant protein vaccines (Figure 2, *p_adj_* < 0.05 for all strains as assessed by a Wilcoxon test with a Bonferroni correction), although there was substantial heterogeneity among individuals (Supplemental Figure 2). However, individuals who received the egg-derived vaccine tended to have clearly lower post-vaccination titers to the 5a.2a.1 clade compared to other viral strains (Figure 2, *p* < 0.001, Mann Whitney U test). A similar trend was not apparent for the recombinant protein vaccine recipients. Note that the post-vaccination titers among the individuals who received the egg-derived vaccine were also lower for a pair of non-5a.2a.1 clade strains that, like 5a.2a.1 clade strains, contain a K142R substitution (gold-highlighted strains near the right of Figure 2, *p* < 0.001, Mann Whitney U test comparing post-vaccination titers among egg-derived vaccine recipients for non-5a.2a.1 strains with and without K142R).

**Figure 2.**
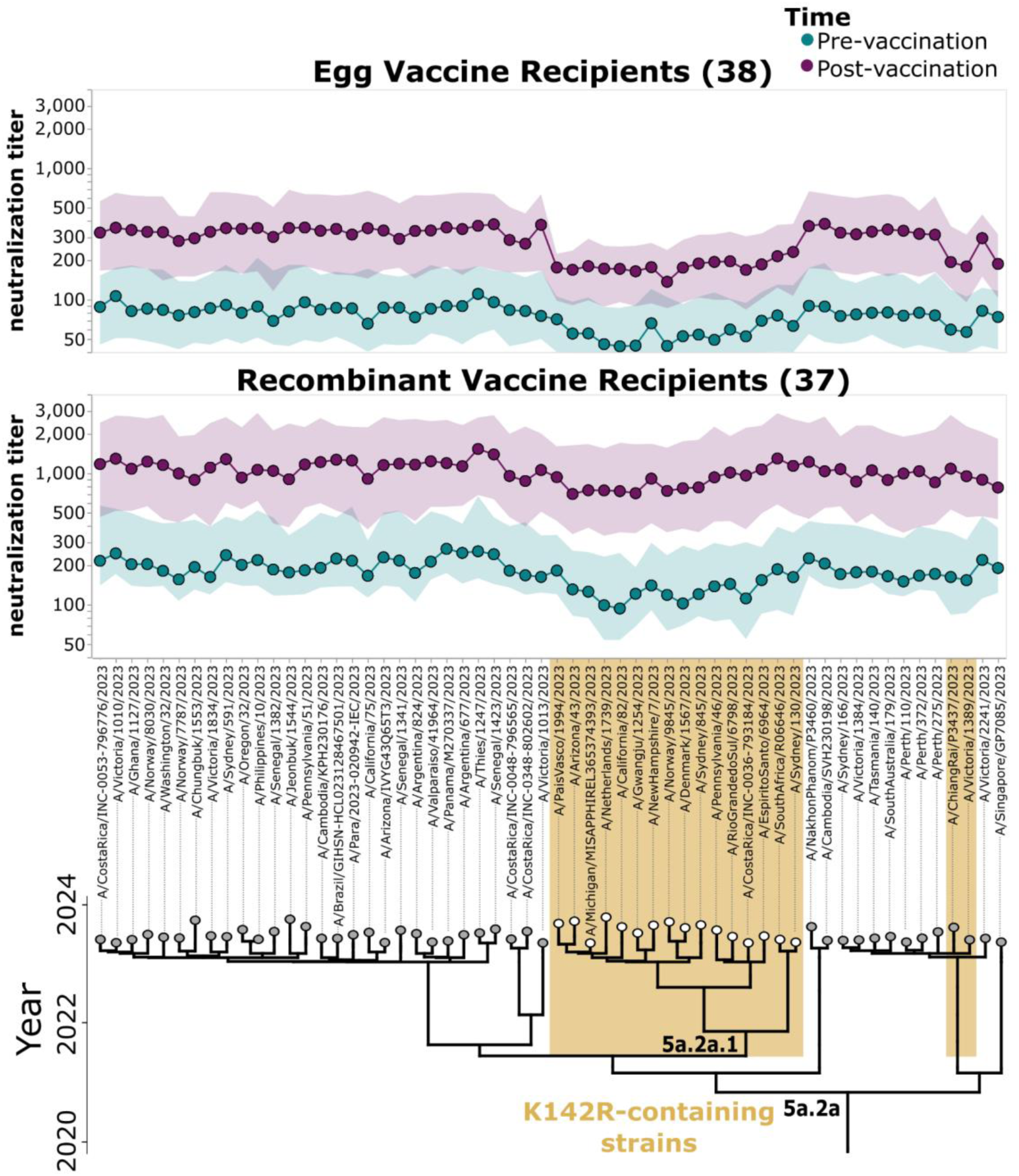
Pre- and post-vaccination titers against a set of H1N1 strains that circulated in 2023/2024 for adults who received egg-derived or recombinant protein vaccines. Neutralization titers against each virus strain pre- and post-vaccination (28-30 days after vaccine receipt) for adults who received either the egg-derived vaccine (top) or the recombinant protein vaccine (bottom) in the 2023/2024 season. Points indicate the median titers, and the shaded areas show the interquartile range. The lower limit of detection given the serum dilutions used was 1:40. Strains are organized by phylogeny on the x-axis, with the 5a.2a clade and the 5a.2a.1 clade (white circles) indicated on the phylogeny. Strains containing K142R mutation are highlighted in gold. Note that the responses and specificity of serum neutralizing antibody titers between different individuals within cohorts is heterogenous, with some individuals showing strain-to-strain variation in titer pre- or post vaccination, and others with broader and more even neutralization potency against all recent strains in the library (see Supplemental Figure 2 for examples).

The pre-vaccination titers are lower against the 5a.2a.1 clade strains than other strains in many individuals within both cohorts (Figure 2). In addition, the median pre-vaccination titers in egg-derived vaccine recipients are approximately ∼2-3 fold lower than those of the recombinant protein vaccine recipients (Figure 2). This fact highlights the caveat that the egg-derived and recombinant protein cohorts are from different studies in different geographic locations, with differences in histories of vaccination with more or less immunogenic vaccines, and so have some differences beyond the vaccine they received. Of note, in temperate regions like the USA there is usually one annual influenza epidemic, while in subtropical regions like Hong Kong there is less regular seasonality and usually two annual influenza epidemics^27,28^, potentially resulting in more exposure to circulating viruses in the recombinant protein vaccine recipients from the Hong Kong study. The lower pre-vaccination titers in the egg-derived vaccine cohort against all strains carry through to after vaccination, with the post-vaccination titers remaining lower against all strains in the egg-derived vaccine cohort compared to the recombinant protein vaccine cohort (Figure 2). Similar trends of lower pre- and post-vaccination titers in the egg-derived vaccine cohort are also observed against prior and current vaccine strains (Supplemental Figure 3). Since there are modest differences in the age ranges of individuals in the egg-derived and recombinant protein vaccine cohorts, we also compared just individuals in the overlapping age ranges (birth years between 1976 and 2001), and found that the trends in titers are extremely similar to the full cohorts (Supplemental Figure 4), suggesting age variation between the cohorts is not responsible for the differences between them (compare Figure 2 and Supplemental Figure 4).

### The recombinant protein vaccine elicits a relatively larger increase in neutralizing antibody titers against strains containing the K142R mutation than the egg-derived vaccine

All three of the HA sites that vary between the egg-derived and recombinant protein vaccine strains, including site 142, are within or adjacent to the Ca2 epitope on the HA head (Figure 3A) ^15^. To assess the effect of each of these mutations individually on neutralization by egg-derived vaccine and recombinant protein vaccine recipients, we examined the fold-change in titer with vaccination against both the egg-derived vaccine strain and cell-vaccine strain used in the recombinant protein vaccine, as well as two circulating strains that represented intermediates between these vaccine strains (Figure 3B). The other two mutations in addition to R142K that distinguish the egg-derived vaccine strain from the recombinant protein vaccine strain are T216A (which varies among strains within the 5a.2a.1 clade) and the common egg-adaptive mutation Q223R^19^. The recombinant protein vaccine elicited a significantly larger titer increase than the egg-derived vaccine against strains that contained the matched identity of R at site 142 (Figure 3B, *p_adj_*< 0.05 as assessed by a Mann-Whitney U test with a Benjamini-Hochberg correction), suggesting that the R142K reversion in the egg-derived vaccine is a major contributor to the strain-specific differences in titers elicited by these vaccines. Titers against the egg vaccine strain (A/Victoria/4897/2022) with and without egg-adaptive mutations (R142K, Q223R) were also assessed by hemagglutinin inhibition assay (HAI) to validate the observation of higher titers against the egg-adapted vaccine strain (IVR-238) post-vaccination in egg-vaccine recipients (Supplemental Figure 5A). HAI measurements were well correlated with sequencing-based neutralization assay titers (Supplemental Figure 5B).

**Figure 3.**
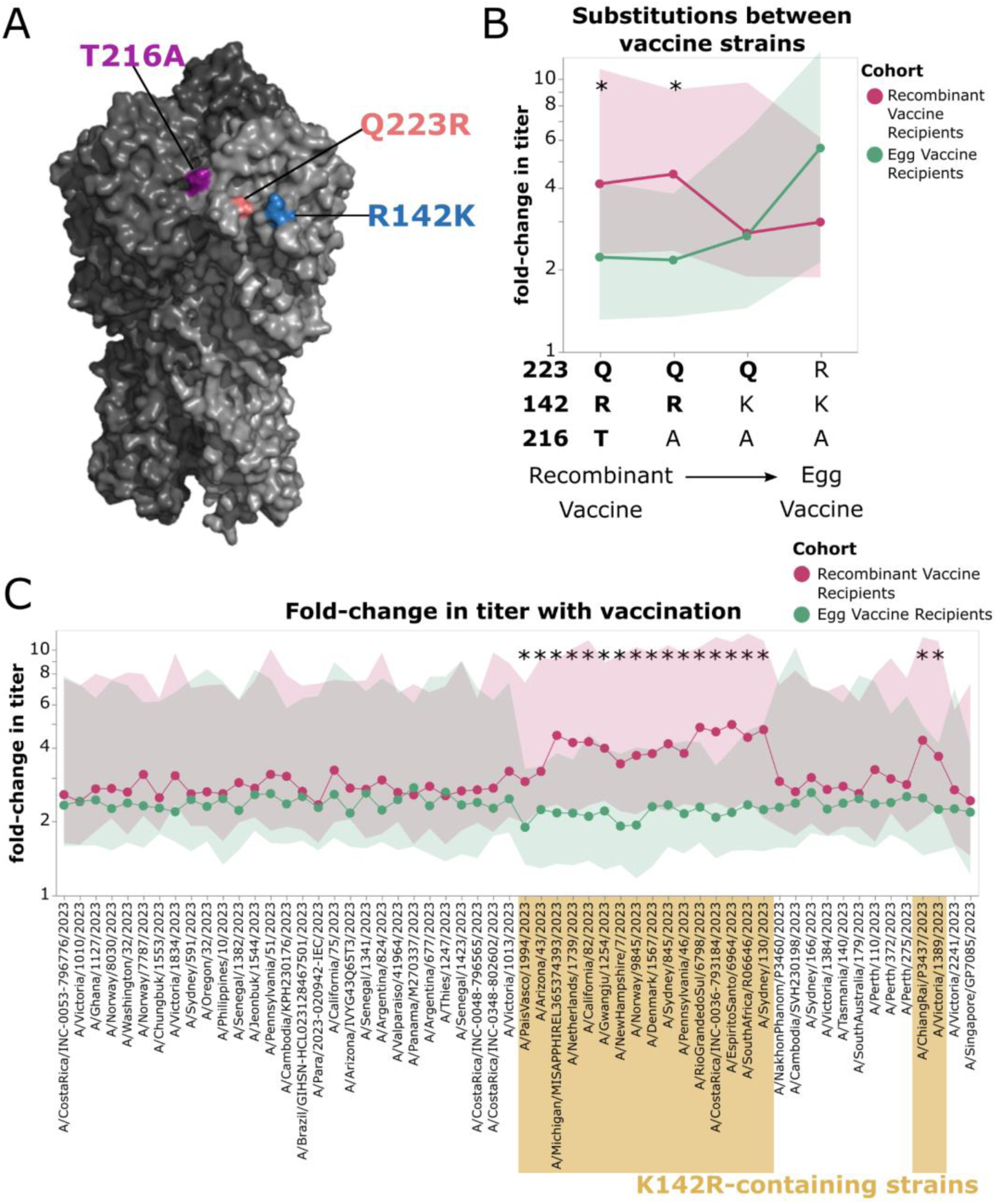
Vaccination-induced change in titers against strains containing K142R differs between egg-derived and recombinant protein vaccine recipients. A) Structure showing the HA amino acid differences between the recombinant protein vaccine strain and the high-growth reassortant used for the egg-derived vaccine (IVR-238) on the HA head (PDB: 6xgc)^15^. B) Titers against the recombinant protein vaccine strain, the egg-vaccine strain, and several other strains that differ only at sites that distinguish the two vaccine strains (in order from left to right: A/Wisconsin/67/2022, A/Michigan/MISAPPHIREL365374393/2023, A/Oregon/Flu-OHSU-241140095/2023, A/Victoria/4897/2022_IVR238). Points and bold lines indicate the median fold-change in titer after vaccination for all sera in that cohort for that strain, and shaded regions show the interquartile range. Strains with a significant difference in the fold-change (p_adj_ < 0.05, after Benjamini-Hochberg correction) between cohorts as assessed by Mann-Whitney U test are indicated with an asterisk. C) Fold-change in titer after vaccination, as measured against the full set of 2023/2024 H1N1 strains. Points and bold lines represent the median fold-change in titer across all sera in that cohort for that strain, and the shaded region shows the interquartile range. Strains with a significant difference in fold-change between groups as assessed by Mann-Whitney U test are indicated with an asterisk (p_adj_ < 0.05, after Benjamini-Hochberg correction). Strains are organized phylogenetically on the x-axis. Strains containing K142R are indicated in gold.

A more comprehensive analysis of the fold-increase in titer across all of the 2023/2024 strains reinforces the idea that the main difference is that the recombinant protein vaccine elicits a larger titer increase against K142R-containing strains compared to the egg-derived vaccine (Figure 3C, which plots the same underlying data as Figure 2 in the form of fold changes). For most strains, both vaccines lead to a similar titer increase of ∼2-3 fold (Figure 3C). But for the K142R-containing strains, which mostly are derived from the 5a.2a.1 clade the vaccine was selected to target, the recombinant protein vaccine elicits a significantly larger titer increase than the egg-derived vaccine (Figure 3C, *p_adj_* < 0.05 as assessed by a Mann-Whitney U test with a Benjamini-Hochberg correction).

### Within vaccine type, age but not prior vaccination, are associated with relatively lower post-vaccination neutralization titers against K142R-containing strains

We next examined whether individuals in certain age groups tended to have lower titers against the K142R-containing 5a.2a.1 clade strains (Figure 4). Prior to vaccination, titers to K142R-containing strains tended to be lower than those to other strains in several individuals across all age groups in both cohorts. Post-vaccination, younger adults (born: 1980-2001) who received the egg-derived vaccine had a stronger trend towards relatively lower (compared to other strains) post-vaccination titers against 5a.2a.1 clade strains and other K142R-containing strains post-vaccination than older individuals (born: 1949-1979) in this cohort. The youngest age-cohort (born: 1990-2001) within the recombinant protein vaccine recipients also exhibits slightly lower relative titers against a subset of 5a.2a.1 clade strains post-vaccination, though this trend is much weaker than for the egg-derived vaccine cohort, and does not appear to be associated with K142R-specificity, as we do not also see lower median titers against non-5a.2a.1 clade K142R-containing strains for this group. Given the small sample sizes within each age group, we are likely underpowered to fully understand the relationship between clade-specificity of titers and age, and other factors, such as geographic location, which could impact historical exposure histories^29^, could also play a role.

**Figure 4.**
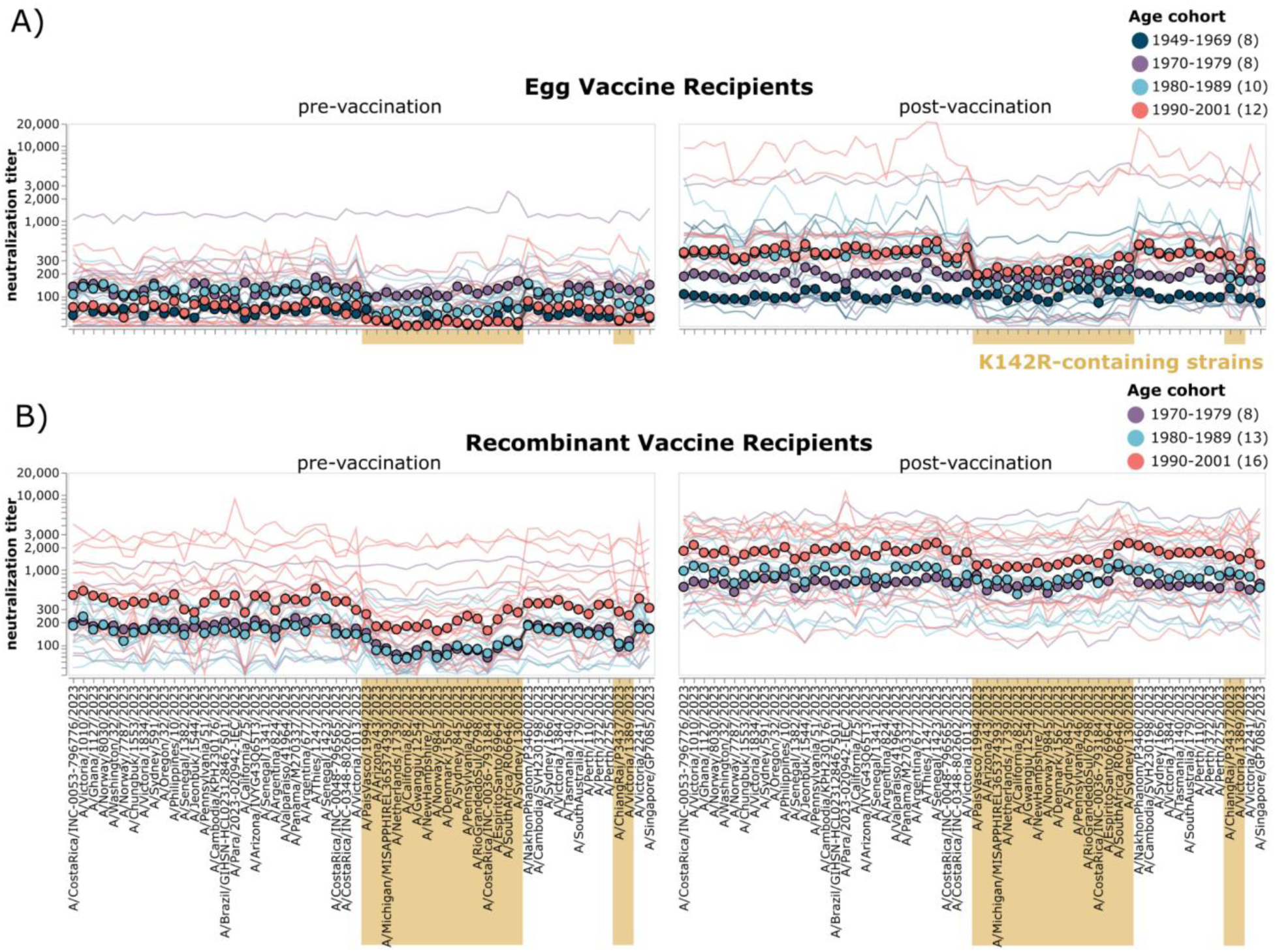
Age as well as vaccine type are associated with relatively lower titers to K142R-containing strains. A) Neutralization titers for pre- and post-vaccination sera from egg-derived vaccine recipients and B) recombinant protein vaccine recipients. Thin lines show neutralization titers against all strains for a single individual, with lines colored by age group. Points represent the median neutralization titer across all sera in that age group for that strain. Strains are organized phylogenetically on the x-axis. Strains containing K142R are highlighted in gold.

Prior vaccination in the two seasons preceding 2023/2024 does not obviously relate to the relatively lower titers to K142R-containing strains compared to other strains (Supplemental Figure 6). In the egg-derived vaccine cohort, both individuals who were vaccinated in both seasons prior to 2023/2024 and individuals who were only vaccinated in one or neither of those seasons have relatively lower titers against K142R-containing strains post-vaccination (Supplemental Figure 6). The absence of an obvious effect of vaccination history on the relative titers to K142R strains is consistent with the fact that prior to 2023/2024 both the egg-derived and recombinant protein vaccines had a lysine at position 142. However, both the egg-vaccine recipient and recombinant protein vaccine recipient cohorts are primarily comprised of individuals who received vaccines in both prior seasons. Therefore, our study may not be well powered to assess the effect of prior vaccination on clade-specific neutralization titers.

### Additional mutations in H1N1 strains in the 2023/2024 season that affect neutralization titers

To assess if additional mutations present in H1N1 strains in 2023/2024 affect neutralization titers, we normalized the titers for each serum to its median titer against all these strains, and looked for strains with higher or lower titers (Figure 5). The mutation among these strains that most affected neutralization was K142R. Mutations at another Ca2 epitope site, P137S/L, also affected titers for multiple individuals in both groups. Both P137S and K142R are present in the 5a.2a.1 clade, and P137S was present in both the recombinant protein and egg-derived vaccine strains. Globally, strains containing K142R and P137S became predominant by June 2025 (Supplemental Figure 7). Several additional mutations in 2023/2024 strains (e.g., A141E, K156R, and K169Q) also have relatively lower neutralization by a handful of sera (Figure 5).

**Figure 5.**
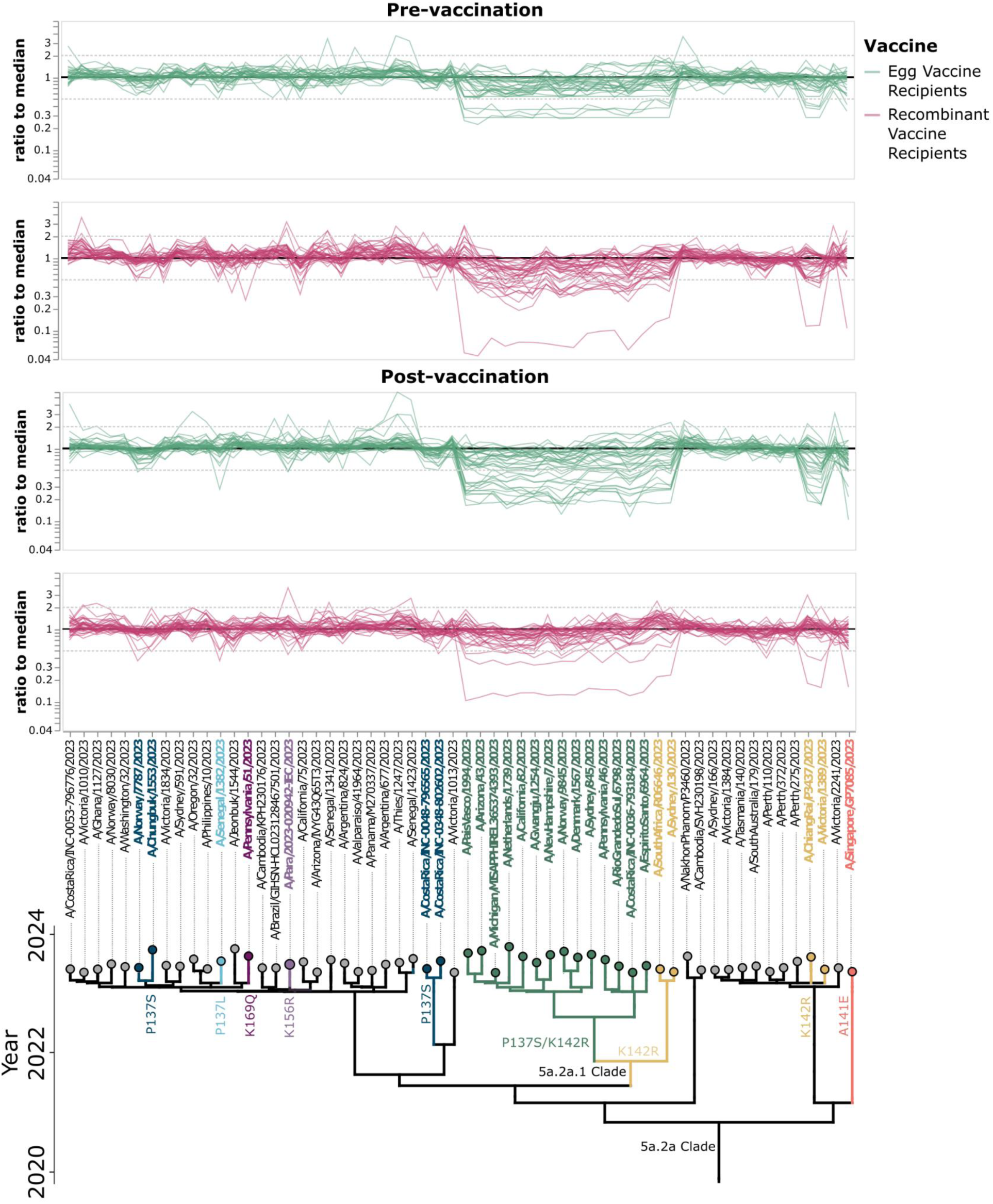
Additional HA mutations occurring in strains circulating in the 2023/2024 season that affect neutralization titers. Thin lines show the ratio of titer for each strain to the median across all the strains for each serum. Strains are organized phylogenetically on the x-axis. Strains or subclades containing mutations that have a greater than two-fold reduction from median titer in at least one serum and contain a mutation in a known epitope site are indicated with coloring, and mutations are annotated on the phylogeny.

## Discussion

We have shown that individuals who received an egg-derived influenza vaccine in the 2023/2024 season had a relatively lower titer boost against a key emerging H1N1 clade compared to other clades than individuals who received a recombinant protein vaccine. Our data suggests that a reversion at site 142 in the high-growth reassortant seed strain used to generate the egg-derived vaccine may have been largely responsible for this effect. Therefore, by making neutralization measurements against many virus strains that circulated in the same influenza season, our study provides empirical data that support the hypothesis^8,9,30,31^ that mutations in the egg-adapted vaccine strain can contribute to clade-specific vaccine differences in vaccine-induced neutralizing titer boosts.

A key limitation of our study is that the cohorts of egg-derived and recombinant protein vaccine recipients were from different studies conducted in different populations. Although our data suggest that the vaccine type received (egg versus recombinant) is the cause of the difference in titers to the K142R-containing 5a.2a.1 clade post-vaccination rather than factors such as age or vaccine history, it remains possible that other unknown factors may confound the relationship between vaccine type used and the observed difference in clade-specific HA responses. For example, it has recently been shown that childhood exposure history may play a role in specificity of antibodies targeting this epitope^29^. To rigorously prove that vaccine type is the determinant of the differences, it would be necessary to have a study in which participants are randomized to receive an egg-derived versus recombinant protein vaccine.

There have been prior observations of mutations in egg-derived influenza vaccine strains altering antigenicity or immunogenicity^32–36^, and potentially resulting in reduced vaccine effectiveness^8,36,37^. However, the majority of influenza vaccines are still produced from viral antigens that have been propagated in eggs, largely because of lower cost and extensive existing infrastructure to make egg-derived vaccines^38^. Understanding how mutations in egg-derived vaccines can impact response to vaccination is critical to a rational approach for developing more effective influenza vaccines.

## Acknowledgements

This work was supported in part by the NIH/NIAID award R01AI165821 to T.B. and J.D.B., U01AI153700 to S.C. and B.J.C., and contract 75N93021C00015 to J.D.B., S.E.H., S.C., B.J.C., and T.B. J.D.B. and T.B. are Investigators of the Howard Hughes Medical Institute. This research was supported by the Genomics & Bioinformatics Shared Resource (RRID:SCR_022606) of the Fred Hutch/University of Washington/Seattle Children’s Cancer Consortium (P30CA015704), and by Fred Hutch Scientific Computing (NIH grants S10-OD-020069 and S10-OD-028685).

We gratefully acknowledge all data contributors for the sequences downloaded from GISAID, including the authors and their originating laboratories responsible for obtaining the specimens, and their submitting laboratories for generating the genetic sequence and metadata and sharing via the GISAID Initiative, on which part of this research is based. A list of laboratories contributing sequences for strains used in this work is provided at https://doi.org/10.55876/gis8.251201rp.

This manuscript is the result of funding in whole or in part by the National Institutes of Health (NIH). It is subject to the NIH Public Access Policy. Through acceptance of this federal funding, NIH has been given a right to make this manuscript publicly available in PubMed Central upon the Official Date of Publication, as defined by NIH.

## Methods

### Data and code availability

All data and code are publicly available at the following links:

- Analysis of the sequencing-based neutralization assays: https://github.com/jbloomlab/flu_seqneut_pdmH1N1_2023-2024_VaccinatedCohorts
- Computational pipeline for analyzing sequencing-based neutralization assays: https://github.com/jbloomlab/seqneut-pipeline
- All measured neutralization titers in CSV format:

- For the Pennsylvania-based adult egg-derived vaccine cohort: https://github.com/jbloomlab/flu_seqneut_pdmH1N1_2023-2024_VaccinatedCohorts/blob/main/results/aggregated_titers/titers_PennVaccineCohort.csv
- For the Hong-Kong-based adult recombinant protein vaccine cohort: https://github.com/jbloomlab/flu_seqneut_pdmH1N1_2023-2024_VaccinatedCohorts/blob/main/results/aggregated_titers/titers_DRIVE.csv
- Interactive visualizations of all neutralization titers: https://jbloomlab.github.io/flu_seqneut_pdmH1N1_2023-2024_VaccinatedCohorts/
- The H1N1 strains in the library listed in CSV format: https://github.com/jbloomlab/flu_seqneut_pdmH1N1_2023-2024_VaccinatedCohorts/blob/update_repostructure/data/viral_libraries/H1N1library_2023-2024_barcode_to_strain.csv

A permanent Zenodo Archive of the GitHub repository containing all of the data, results, and computer is available at DOI: https://doi.org/10.5281/zenodo.19631554.

### Human sera

Human sera from an observational adult vaccine cohort based in Philadelphia were taken from adults (ages 22-74) on the day of vaccination and 28 days post vaccination with a 2023/2024 Northern Hemisphere egg-derived vaccine (FluLaval quadrivalent influenza virus vaccine from GlaxoSmithKline) between October-December 2023. These participants also provided information on self-reported influenza vaccination history in the past two years (2021/22 and 2022/23) (Table 1). This study was approved by the Institutional Review Board of the University of Pennsylvania under protocol number 849398.

Human sera from an interventional study based in Hong Kong were selected from a randomized vaccine trial of repeated annual vaccination of Flublok done in Hong Kong in 2020–2025 (DRIVE-1 study)^39^. The DRIVE-I study (ClinicalTrials.gov: NCT04576377) is a randomized placebo-controlled trial of repeated annual influenza vaccination in adults 18–45 years of age at enrollment (in 2020/21). For the present study, we selected samples provided by participants from year 4 (i.e. 2023/24), with a similar distribution of vaccinated years as compared to the egg-derived vaccine cohort (Table 1). The vaccine used, Flublok, is a quadrivalent recombinant-HA vaccine produced in insect cells, including 45 mcg HA of each of the four included strains as recommended by the World Health Organization for the northern hemisphere in the 2023–2024 season. Sera used in this study was collected immediately before administration of vaccine/placebo, and again at a follow-up visit approximately 30 days after vaccination. The study protocol was approved by the Institutional Review Boards of the University of Hong Kong (ref: UW19-551) and of the University of Chicago Biological Sciences Division (ref: IRB20-0217). Written informed consent was obtained from all participants.

Prior to use in sequencing-based neutralization assays, all sera were treated with receptor-destroying enzyme and heat-inactivated, as described previously^40^. Briefly, one vial of lyophilized receptor-destroying enzyme II (Seikan) was resuspended in 20 mL PBS and passed through a 0.22 uM filter. Then, each serum sample was diluted 1:4 in the resuspended receptor-destroying enzyme solution, and incubated at 37°C for 2.5 hours and then 55°C for 30 minutes. Sera were then used immediately or stored at -80°C until use.

### Selection of strains with HAs to include in the H1N1 library

To identify representative circulating strains, we used H1N1 Nextstrain^41,42^ builds available in November 2023. These trees are subset by Nextstrain-defined clade, subclade and derived haplotype, where a derived haplotype is a more fine-grained level of genetic subdivision than subclade, and is defined as a subset of strains belonging to the same subclade that each share additional amino acid mutation(s) and have achieved some threshold of child strains. We filtered all derived haplotypes by collection date, retaining haplotypes with a strain sequenced within the 6-months prior to library design in November of 2023. For each of the derived haplotypes we selected a naturally-occurring HA sequence with the lowest divergence from the derived haplotype consensus sequence. This yielded 58 strains, representing the diversity of H1N1 HA strains circulating globally in 2023. Additionally, the past decade of cell-passaged H1N1 vaccine strains were added (A/California/07/2009, A/Michigan/45/2015. A/Brisbane/02/2018, A/Hawaii/70/2019, A/Wisconsin/588/2019, and A/Wisconsin/67/2022) as well the recent egg-based vaccine (A/Victoria/4897/2022_IVR238), and one strain similar to A/Victoria/4897/2022_IVR238 but lacking the egg-associated mutation Q223R (A/Oregon/Flu-OHSU-241140095/2023), constituting an additional 8 strains.

### Design of H1N1 barcoded HA genomic segments

To insert a unique barcode into the HA genomic segment without substantially disrupting vRNA packaging, we designed a construct with a duplicated packaging signal at the 5′ end of the negative-sense HA vRNA, as described previously^11,43,44^.

### Generation of H1N1 barcoded viruses included in library

Unidirectional reverse genetics plasmids encoding all barcoded variants of the same HA sequence were pooled at equal concentrations and used to generate influenza viruses containing unique barcodes by reverse genetics, with the WSN pHW18* series of plasmids for all seven non-HA viral genes ^45^. The non-HA genes from A/WSN/1933 (H1N1) were used because this laboratory-adapted strain consistently produces high titer viral stocks by reverse genetics, and its extensive laboratory adaptation is considered to have substantially reduced its risk to humans. To produce these viral stocks, briefly, 5e5 293T cells and 5e4 MDCK-SIAT1-TMPRSS2 cells were plated in a 6-well dish in D10 media (DMEM supplemented with 10% heat-inactivated fetal bovine serum, 2 mM L-glutamine, 100 U per mL penicillin, and 100 µg per mL streptomycin). Approximately 24 hours after plating cells, a master mix containing 250 ng/well of each of the 7 internal reverse genetics plasmids for WSN virus and the pooled barcoded HA plasmids was prepared in 100 μL DMEM with 3 μL BioT reagent. This was incubated for 15 minutes at room temperature and then added dropwise to the plate. At 20-24 hours post-transfection, media was removed, and cells were washed once with phosphate-buffered saline (PBS) and 2 mL Influenza Growth Media (Opti-MEM supplemented with 0.1% heat-inactivated FBS, 0.3% bovine serum albumin, 100 µg per mL of calcium chloride, 100 U per mL penicillin, and 100 µg per mL streptomycin) was added to cells. Then at approximately 65 hours post-transfection, cell supernatants containing the barcoded influenza viruses were collected and centrifuged for 4 min at 845 × g to remove cell debris. Aliquots of clarified viral supernatants were then frozen at −80°C for storage. To expand viruses to high titer, all virus pools generated by reverse genetics were passaged once in MDCK-SIAT1-TMPRSS2 cells. For this, MDCK-SIAT1-TMPRSS2 cells were seeded at a density of 4e5 cells/well in a six-well dish in D10, and at 4 hours after seeding were washed with PBS and 2 mL of Influenza Growth Medium was added to each well. Each well was inoculated with 100 μL of the virus supernatant from reverse genetics and the virus was allowed to grow in the cells for ∼40 hours. Cell supernatants were then collected and clarified by centrifugation at 845 × g for 4 minutes, and aliquots of clarified viral supernatants were stored at −80°C for use in the sequencing-based neutralization assay.

### Pooling of virus library strains

To determine the ratios at which to pool barcoded library strains, we used the same approach as previously described^11^. Briefly, an equal volume pool of library strains was made and this was serially diluted in a 96-well plate. Then, 50,000 MDCK-SIAT1 cells were added to each well of the plate. Then, approximately 16 hours after infection, viral barcodes were prepared for sequencing as described below. Sequencing counts for each barcoded strain were used to determine the relative contribution of each HA strain to the virus pool. We then calculated the relative amount of each viral strain so that each strain’s barcodes should correspond roughly to 1/64 of the sequencing counts for the total pool, and re-pooled each of the strains based on these data. This re-pooled library was again serially diluted and used to infect MDCK-SIAT1 cells added at 5e4 cells per well to 96-well plate, and viral barcodes were sequenced and used to verify each strain (and the barcodes corresponding to each strain) was present at relatively equal amounts. These data were also used to determine the infection conditions under which the amount of vRNA from the infecting library linearly corresponded with the amount of library added to each well. We chose to use a dilution factor of 1:22.5 for this particular virus library. While 50,000 cells/well were used in this initial test, as we had seen in other experiments that increasing cell number expanded the linear range at which this assay could be conducted, we chose to use 100,000 MDCK-SIAT1 cells/well in subsequent assays. MDCK-SIAT1 cells were used because they lack TMPRSS2 (which cleaves HA in producing cells and is important for producing infectious virions). These cells should have only limited secondary viral replication.

In the initial pool, we had attempted to include a A/Victoria/4897/2022-like strain that contained K142R, however, this strain was not detected in the initial pool, indicating that we were unable to rescue this virus by reverse genetics. Given this, we rescued and passaged two additional strains, according to the method described above (A/Victoria/4897/2022_IVR238 and A/Oregon/Flu-OHSU-241140095/2023). These strains were added to allow us to better understand both the effects of the mutations that differed between the cell and egg vaccines. A stock of these two strains was prepared and mixed with the pooled library prior to each assay such that these strains were represented at a similar concentration as other variants in the library.

### Sequencing-based neutralization assay

The experimental approach used in this assay is nearly identical to that outlined previously^11^, incorporating a few modifications for increased library size, which have previously been described^12^. An updated protocol including these modifications has been made available (dx.doi.org/10.17504/protocols.io.ewov1962klr2/v1). The setup for the sequencing-based neutralization assay protocol is performed similarly to other 96-well plate-based neutralization assays. First, human sera that had been pre-treated with receptor-destroying enzyme was serially diluted 2-fold across the plate, using an initial dilution of 1:40 (accounting for the 1:4 dilution that was used during the receptor-destroying enzyme treatment as described above, and initially prepared at 2x desired concentration, to account for the volume of virus library that would be added to each well). This dilution series was performed in Influenza Growth Media and comprised 11 columns of the 96-well plate, with a final volume of 50 uL diluted serum in each well. The final column of the 96-well plate was used for a no-serum control, and contained only 50 uL of Influenza Growth Media. A 50 uL volume of diluted virus library was then added to each well, and virus-serum mixtures were incubated at 37°C with 5% CO2 for 1 hour. Following this incubation, 1e5 MDCK-SIAT1 cells were added to each well in a total of 50 uL of Influenza Growth Media and then incubated for approximately 16 hours. At approximately 16 hours post-infection, cells were lysed and barcodes were sequenced as described below.

### RNA extraction and barcode sequencing

Our sequencing protocol was performed similarly to that described previously^11,12^. Briefly, supernatant was removed and cells were washed once with 150 uL of phosphate buffered saline before each well was lysed with 50 uL of iScript RT-qPCR Sample Preparation Reagent (BioRad) containing the barcoded RNA spike-in (which was generated, purified and quantified as described in Loes et al.^11^) at 2 pM per well. Sequencing of library pooling plates was done exactly as previously described^11^, but sequencing for neutralization assay plates was performed with the modification described in Kikawa et al^12^.The lysis reaction proceeded for 5 minutes before lysate was transferred off cells and into a new 96-well plate. Then, 1 uL of this lysate was added to 9 uL cDNA synthesis mastermix for all wells using the iScript Select cDNA Synthesis Kit (BioRad) according to the manufacturer’s instructions, using a gene-specific primer 5’-CCTACAATGTCGGATTTGTATTTAATAG-3’.

The cDNA produced was amplified using two rounds of PCR. In the first round, either the Round 1 sequencing forward primer described in Loes et al.^11^ (5′-*GTGACTGGAGTTCAGACGTGTGCTCTTCCGATCTCTCCCTACAATGTCGGATTTGTATTTAATAG-*3′) was used, or the following forward primers were used to incorporate a 6-bp plate-based index, as described in Kikawa et al. ^12^. These forward primers were paired with the same reverse primer described in Loes et al.^11^ (5′-AGCAAAAGCAGGGGAAAATAAAAACAACC-3′). The set of forward primers was as follows: 5’-GTGACTGGAGTTCAGACGTGTGCTCTTCCGATCTgctacaCCTACAATGTCGGATTTGTATTTAA TAG-3’, 5’-GTGACTGGAGTTCAGACGTGTGCTCTTCCGATCTatcgatCCTACAATGTCGGATTTGTATTTAAT AG-3’, 5’-GTGACTGGAGTTCAGACGTGTGCTCTTCCGATCTtgacgcCCTACAATGTCGGATTTGTATTTAA TAG-3’, 5’-GTGACTGGAGTTCAGACGTGTGCTCTTCCGATCTcagttgCCTACAATGTCGGATTTGTATTTAAT AG-3’.

By incorporating these plate-based indices, we were able to multiplex different plates using the same Round 2 dual-indexing UDI primers, decreasing sequencing costs. Regardless of which forward primer was used for round 1 PCR, we used 5 uL of cDNA as template in a 50 uL reaction using KOD Polymerase Hot Start 2x Mastermix (Sigma) according to the manufacturer’s instructions. The second round of PCR adding unique dual indexing primers was performed exactly as described previously, and after PCR samples were pooled at equal volume and ran on a 1% agarose gel at 85V for 35 minutes. Resulting bands were extracted and purified using Nucleospin Gel Extraction Kit (Takara) before purification with Ampure XP beads (Beckman Coulter). The purified pool was then quantified using a Qubit dsDNA High Sensitivity Kit (Thermo Scientific). The indexed, purified and quantified DNA was then diluted to 4 nM and submitted for Illumina Sequencing, with a target of at least 5e5-1e6 reads per well. A more detailed PCR protocol is described on <u>protocols.io</u> at dx.doi.org/10.17504/protocols.io.ewov1962klr2/v1.

### Analysis of sequencing-based neutralization data

The analysis of data obtained from sequencing-based neutralization assays was performed using the modular *seqneut-pipeline* v3.1.3 (https://github.com/jbloomlab/seqneut-pipeline) which calculates fraction infectivities from normalized barcode counts (as described in Supplemental Figure 1), and then fits Hill curves to fraction infectivity values using the Python package *neutcurve* (https://github.com/jbloomlab/neutcurve). The processing of these data has been described previously^11^ and in the *seqneut-pipeline* README (https://github.com/jbloomlab/seqneut-pipeline/blob/main/README.md). Neutralization titer data for this study and additional information regarding library composition and design are available at https://github.com/jbloomlab/flu_seqneut_pdmH1N1_2023-2024_VaccinatedCohorts.

A single serum sample (PENN23_y1964_s006) was excluded from cohort level analysis as the timepoints for this sample appeared to have been mixed up during the experimental assays (i.e. titers decreased for several recent strains and vaccine strains post-vaccination). Titers for this individual pre- and post-vaccination are available in titer files. All analysis plots were run with and without this individual’s sample to confirm that inclusion or exclusion did not impact the overall results or significance of any comparisons that are described.

### Statistical tests

All statistical comparisons that were completed are available within the associated project repository (https://github.com/jbloomlab/flu_seqneut_pdmH1N1_2023-2024_VaccinatedCohorts/tree/main/notebooks). Briefly, To assess if individuals in the egg-derived vaccine cohort had lower titers against either the 5a.2a.1 clade compared to other clades or simply the non-5a.2a.1 clade K142R-containing strains compared to other strains, we performed a one-sided Mann-Whitney U Test comparing all titers for this group against first the 5a.2a.1 clade strains vs. all 5a.2a strains, and then against K142R strains in 5a.2a vs strains with an arginine at 142 in 5a.2a clade (A/ChiangRai/P3437/2023, A/Victoria/1389/2023). To assess differences in fold-changes between the egg-derived vaccine recipients and the recombinant vaccine recipients, we also performed a one-sided Mann-Whitney U test comparing the fold-change in titers for each strain between the two groups and applied a Benjamini-Hochberg correction. We found that all strains in 5a.2a.1 or with K142R had significantly different fold-changes between groups after a Benjamini-Hochberg correction. Additionally, no strains with a lysine at position 142 had statistically significant differences in fold-change in titer between the two groups after a Benjamini-Hochberg correction. Finally, to determine whether there were statistically significant differences in neutralization of K142R-containing strains between age cohorts, we calculated the geometric mean titer (GMT) against K142-containing strains and R142-containing strains for each individual, then calculated the difference in GMT. We then performed a one-sided Mann-Whitney U test to assess if there was a statistically significant difference in the GMTs between the individuals from the 1980-2001 cohort compared to older individuals. This was not significant (*p* = 0.0503). We also performed a similar test comparing individuals in the 1990-2001 cohort of recombinant vaccine recipients to older individuals. This was also not significant (*p* = 0.24).

### Hemagglutination-inhibition (HAI) assays

For the hemagglutination-inhibition assays in Supplemental Figure 4, serum samples were first treated with receptor-destroying enzyme (RDE) (Denka Seiken) at 37°C for two hours, followed by heat inactivation at 55°C for 30 minutes. RDE-treated samples were then incubated with 10% (vol/vol) turkey red blood cells (RBCs) (Lampire) and placed at 4°C for one hour. After incubation, tubes were spun down and treated sera were transferred to clean tubes and stored at 4°C until use. Treated samples were serially diluted 2-fold in DPBS in 96-well round bottom plates (Corning), followed by the addition of four agglutinating doses of viruses diluted in DPBS in a total volume of 100 µL. Next, 12.5 µL of 2% (vol/vol) turkey RBCs were added to each well and mixed gently with sera and virus. After one hour of incubation at room temperature, plates were tilted vertically for one minute allowing the formation of teardrop shapes of RBCs and plates were scanned. HAI titers were determined as the inverse of the highest dilution that inhibited agglutination.

## Supplemental Figures

**Supplemental Figure 1.**
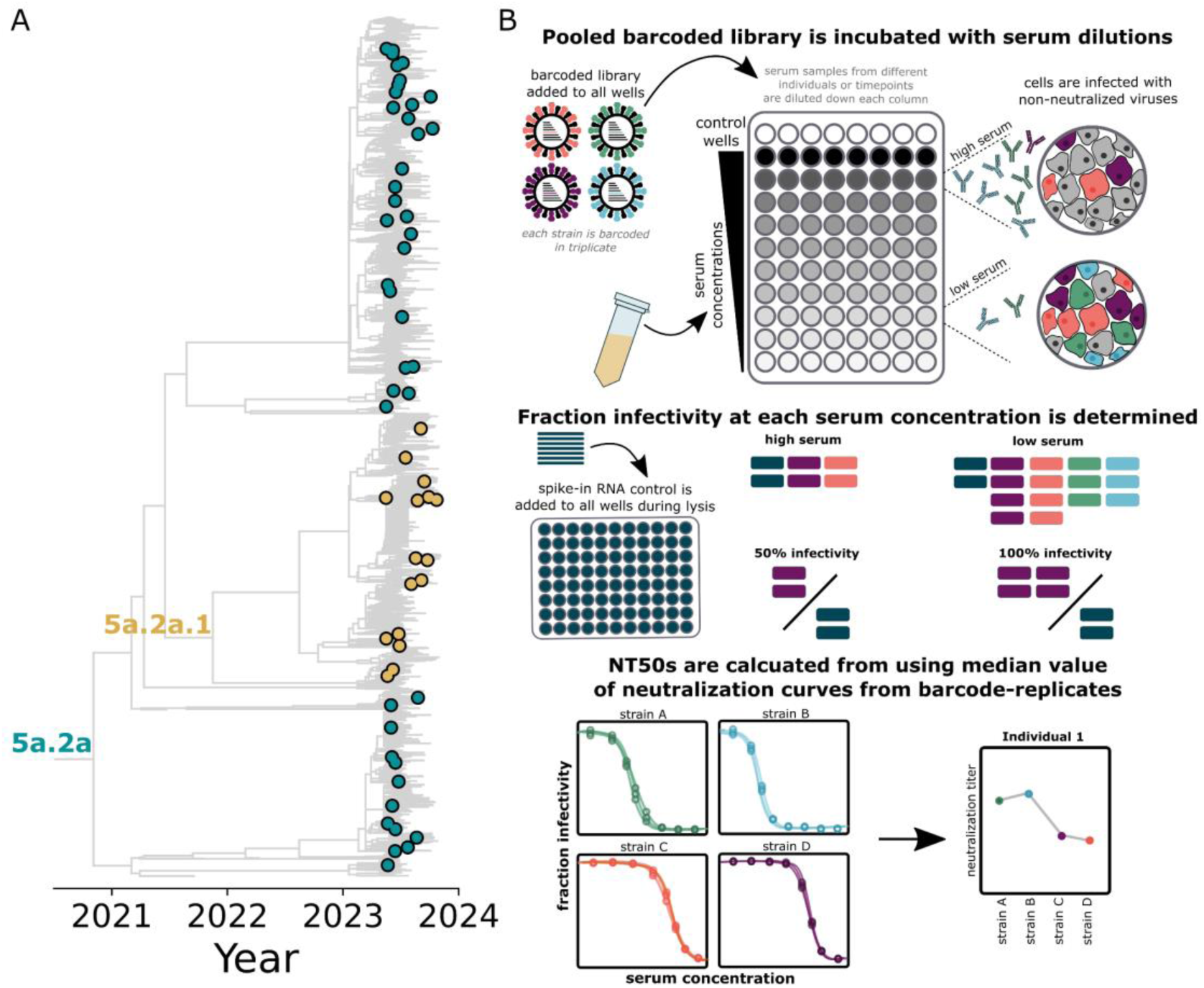
Strains selected for the sequencing-based neutralization assay represent the diversity of H1N1 viruses circulating in the fall of 2023. **A)** Phylogenetic tree of strains selected representing top haplotypes circulating in fall of 2023 from https://nextstrain.org/seasonal-flu/h1n1pdm/ha/6m@2023-11-16. Strains selected for inclusion in the library are indicated with circles. Strains within the 5a.2a.1 clade are labelled in gold and other 5a.2a clade strains are labelled in teal. B) Schematic of the sequencing-based neutralization assays. Briefly, pooled barcoded viruses are incubated with serum dilutions for 1 hr, and then cells are added such that non-neutralized viruses can infect and amplify barcoded HA vRNA segments. At 16 hrs, a barcoded RNA spike-in control is added at the same concentration to each well, such that the fraction infectivity of each barcoded strain can be estimated from the fraction of that barcode to the fraction of the barcoded RNA spike-in control in a given well, compared to the fraction at which that barcode is represented in the no-serum control wells. Curves are then fit to the fraction infectivity measurements to calculate a neutralization titer at 50% infectivity (or NT50).

**Supplemental Figure 2.**
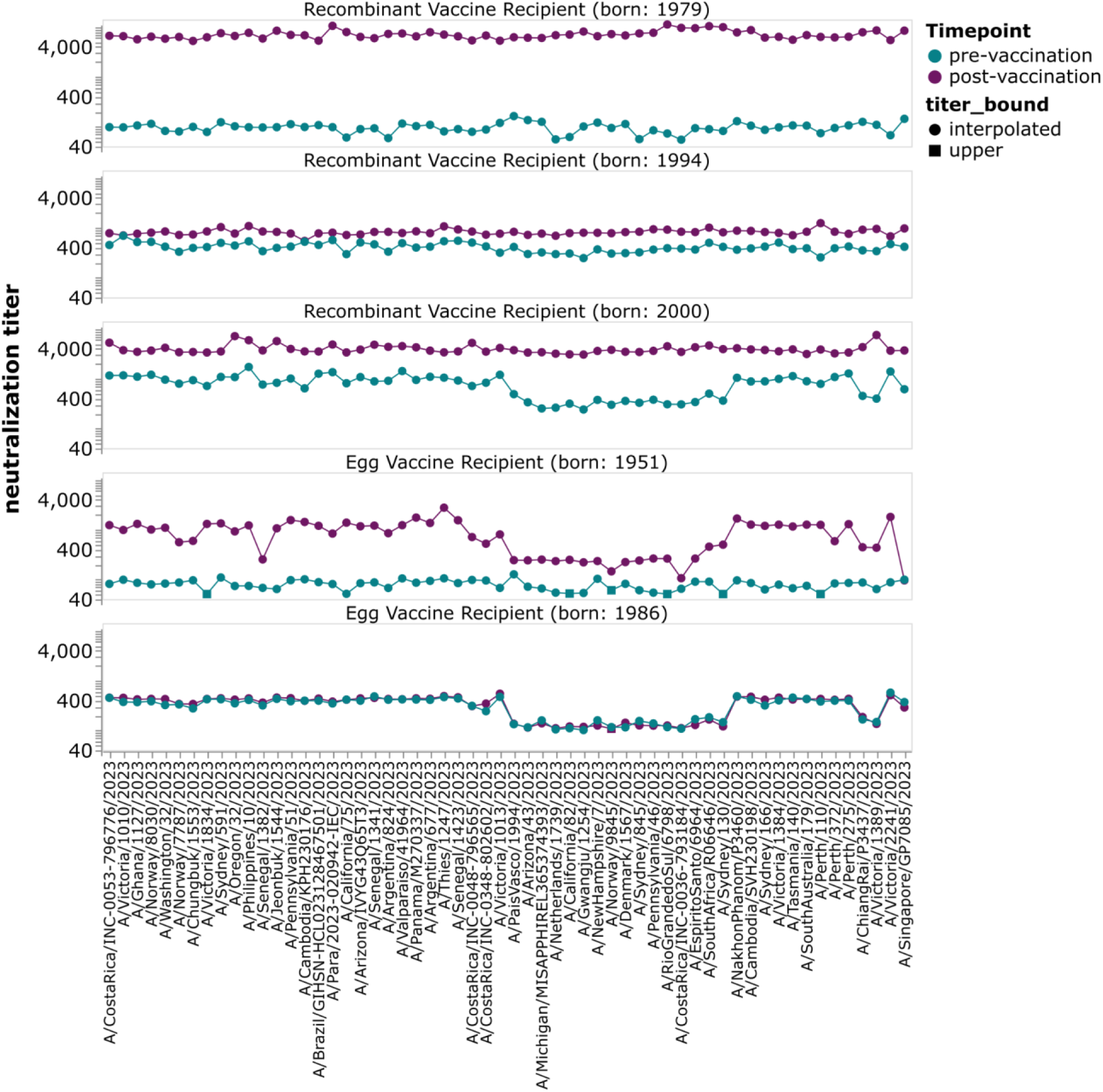
Heterogeneity among individuals in response to vaccination and neutralizing titers to recent strains. Neutralization titers against the library of 2023 H1N1 strains for pre- and post-vaccination sera from some example individuals who received either the recombinant protein or egg-derived vaccine. The points represent the titer against each strain. Strains are grouped phylogenetically along the x-axis. Some calculated titers were outside the range of dilutions tested (titer_bound); NT50s shown as squares, indicated as being at the upper bound, were below the lowest dilution tested.

**Supplemental Figure 3.**
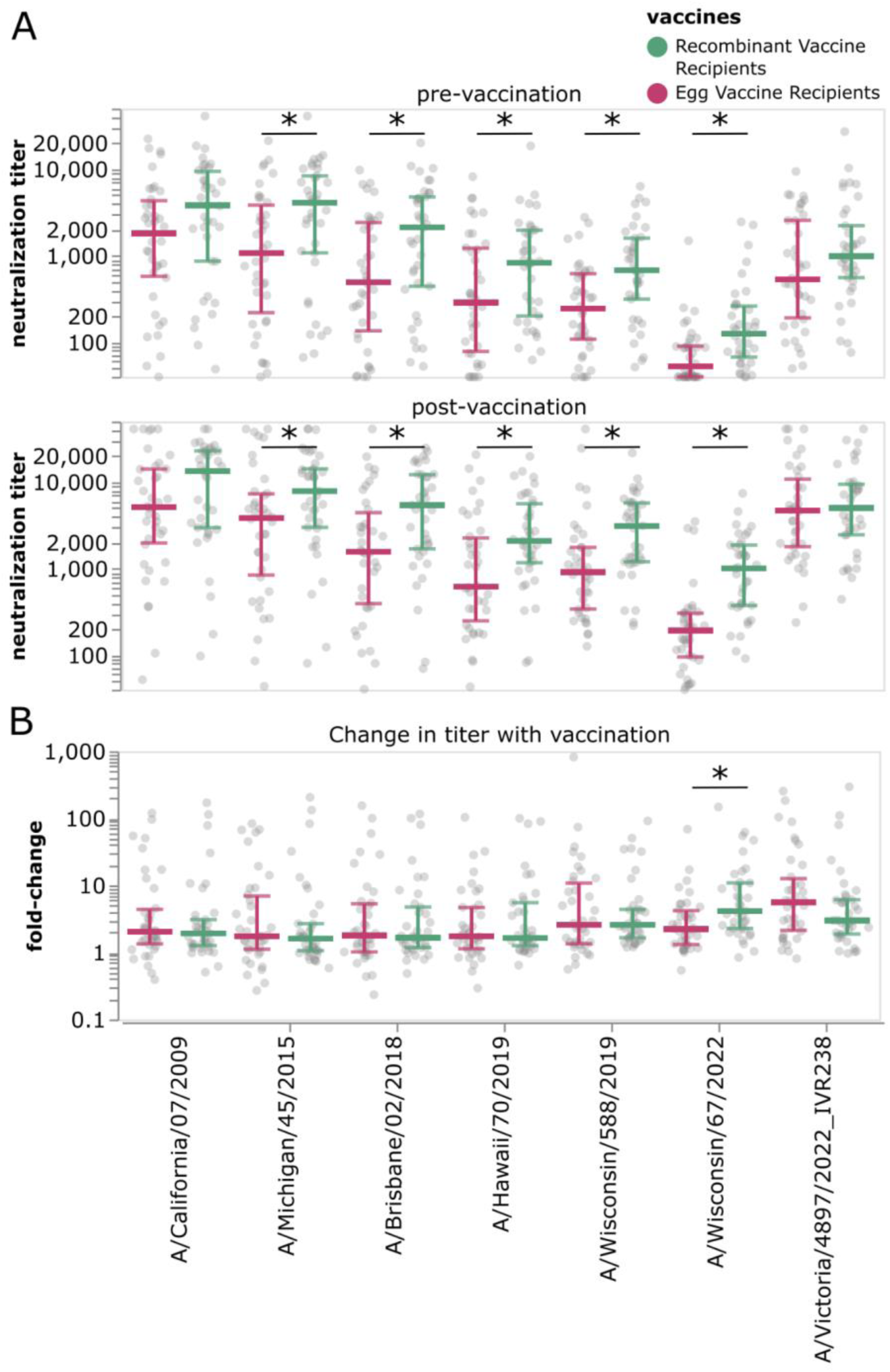
Neutralizing antibody titers to prior vaccine strains. A) Neutralization titers pre- and post-vaccination against previous seasons’ vaccine strains. B) Fold-change in titer against previous vaccine strains after vaccination. Grey points show the measured titer or fold-change for a single individual in each group. Colored horizontal lines represent the median titer for all individuals in a given group against that vaccine strain. Error bars show the interquartile range for that group. Strains with a significant difference in titer or fold-change between groups as assessed by Mann-Whitney U test are indicated with an asterisk.

**Supplemental Figure 4.**
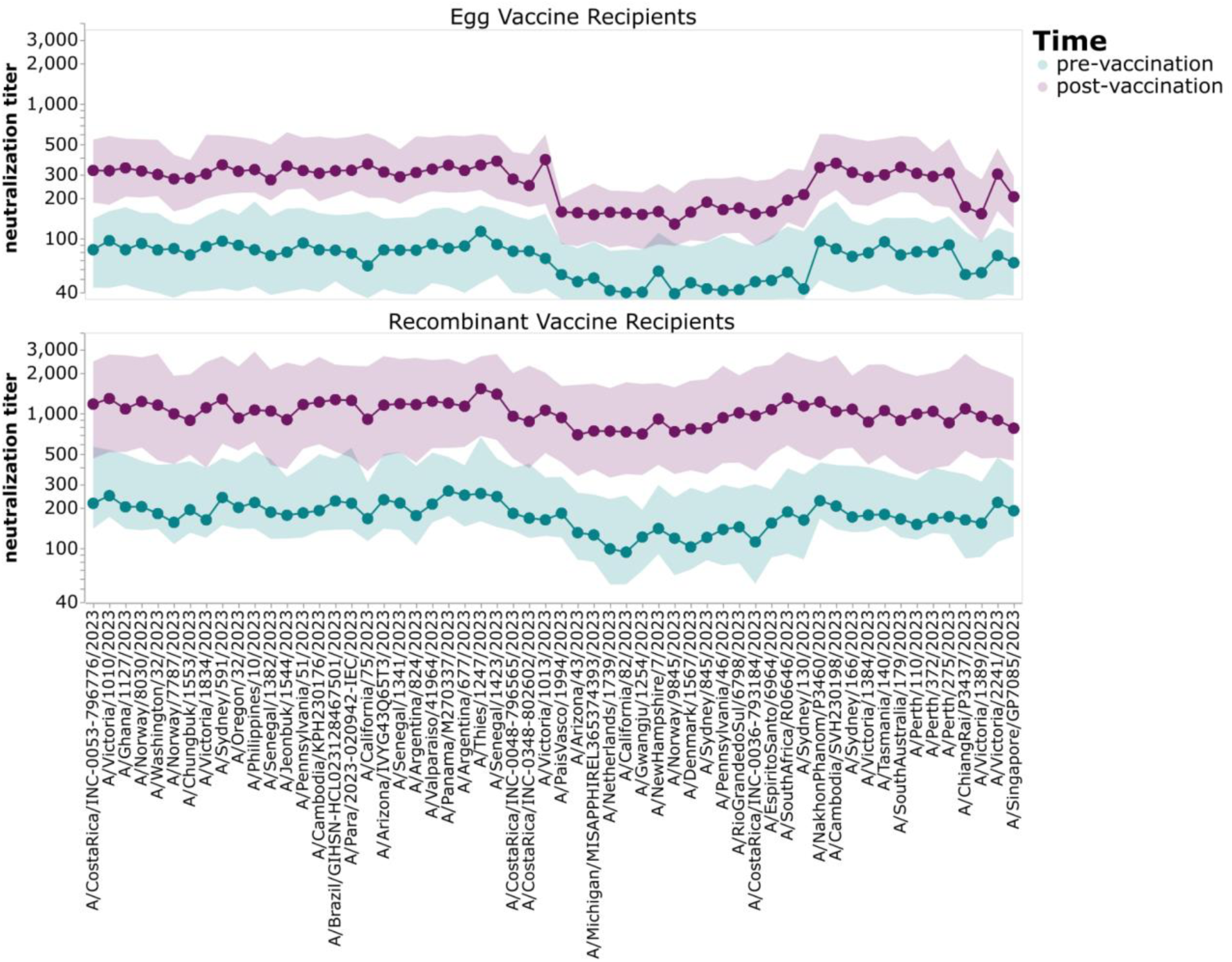
Pre- and post-vaccination titers against a set of H1N1 strains that circulated in 2023/2024 for adults born between 1976 and 2001 who received egg-derived or recombinant protein vaccines. This figure shows data comparable to Figure 2 except it includes only the individuals in the cohorts that were born between 1976 and 2001.

**Supplemental Figure 5.**
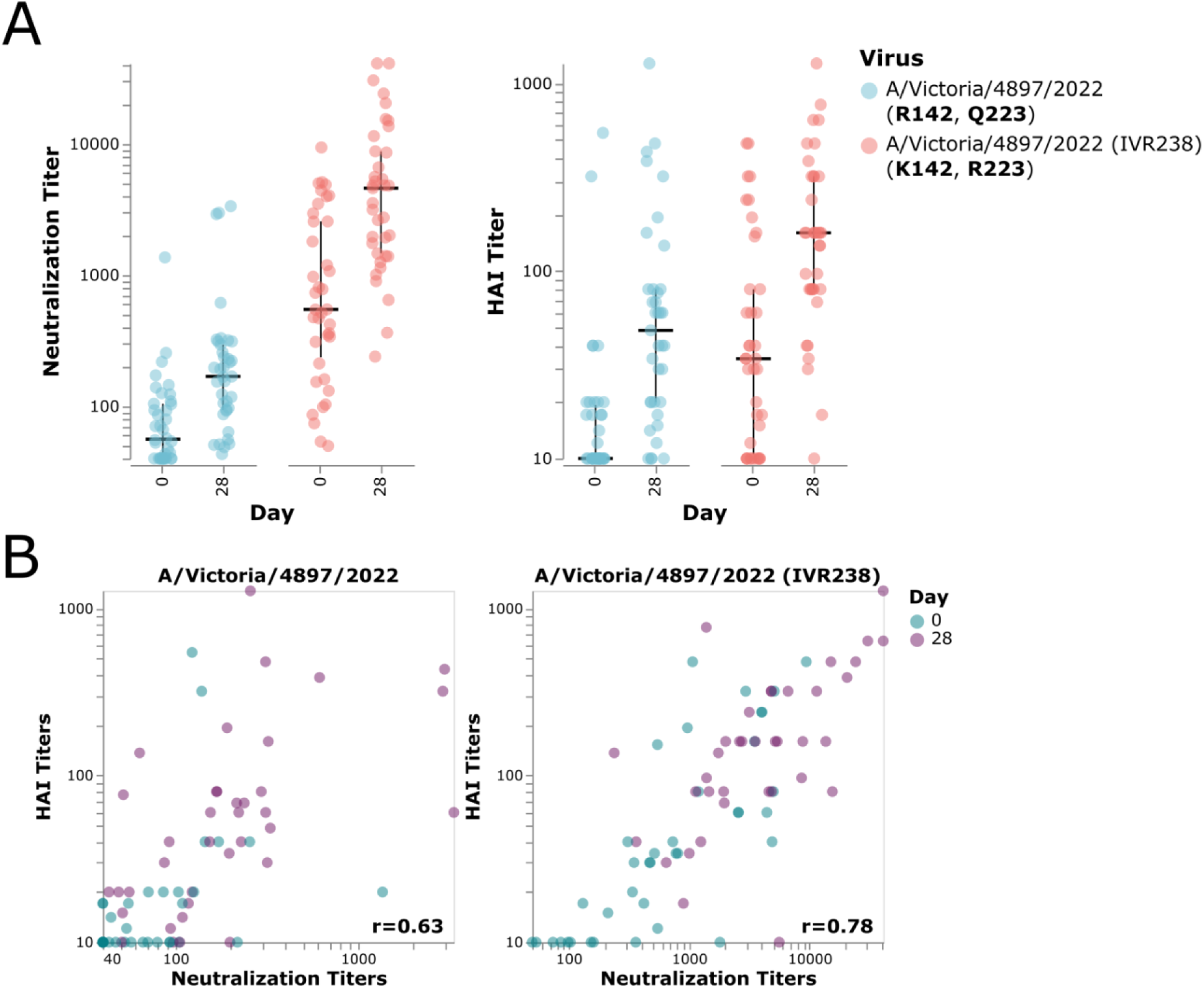
Comparison of hemagglutinin inhibition titers (HAI) and sequencing based-neutralization titers against the A/Victoria/4897/2022 (IVR238) vaccine strain and A/Victoria/4897/2022 strain without egg-adaptive mutations (R142K, Q223R) for individuals who received the egg-derived vaccine. A) Titers collected with sequencing-based neutralization assay and with HAI assay against A/Victoria/4897/2022 and A/Victoria/4897/2022(IVR-238) show similar trends. Points show measurements for each individual against each strain at day 0 or day 28, respectively. Horizontal lines show median values for that group and vertical bars indicate the interquartile range. B) Titers at day 0 and day 28 collected by either HAI or with sequencing-based neutralization assay are well correlated. Points show measurements for each individual against each strain at day 0 or day 28, respectively and are colored by the day sera was collected. Pearson correlations (r) are indicated.

**Supplemental Figure 6.**
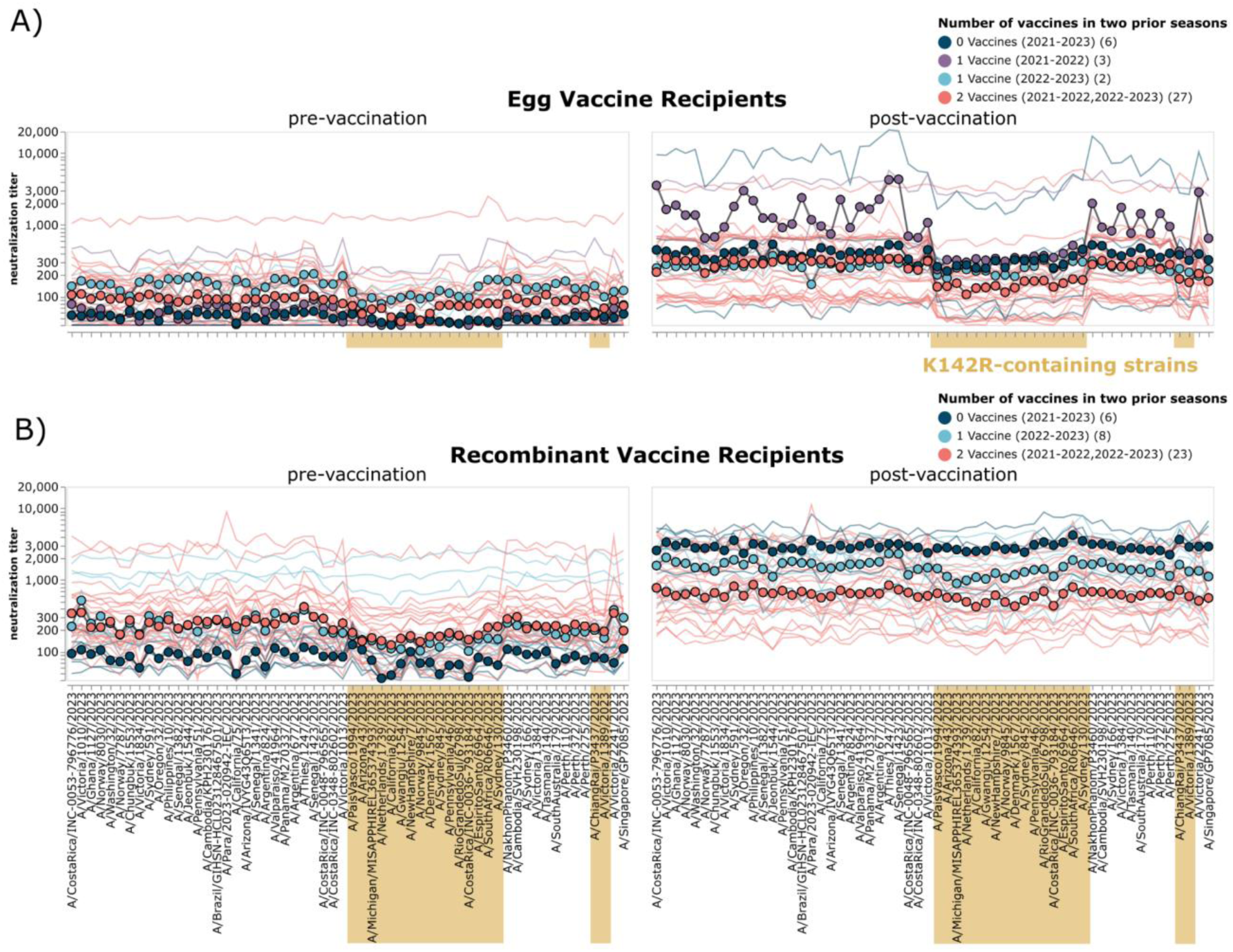
Vaccination history in the two seasons preceding 2023/2024 is not obviously related to the relatively lower titers to K142R-containing strains. A) Neutralization titer profiles for pre-and post-vaccination sera from egg-derived vaccine recipients. B) Neutralization titer profiles for pre- and post-vaccination sera from recombinant vaccine recipients. Thin lines show a neutralization titer profile for a given individual. Each point represents the median neutralization titer across all sera in that group for that strain. Strains are organized phylogenetically on the x-axis. Strains containing K142R are indicated in gold. A shared x-axis is used for both panels and a colored bar below panel A is provided to facilitate easier association with labeled strains. Recombinant vaccine recipient group with 2 vaccines in 2021-2023 seasons includes individuals with and without vaccines in 2020-2021 season. Vaccination status for egg-derived vaccine recipients in 2020-2021 season is unknown.

**Supplemental Figure 7.**
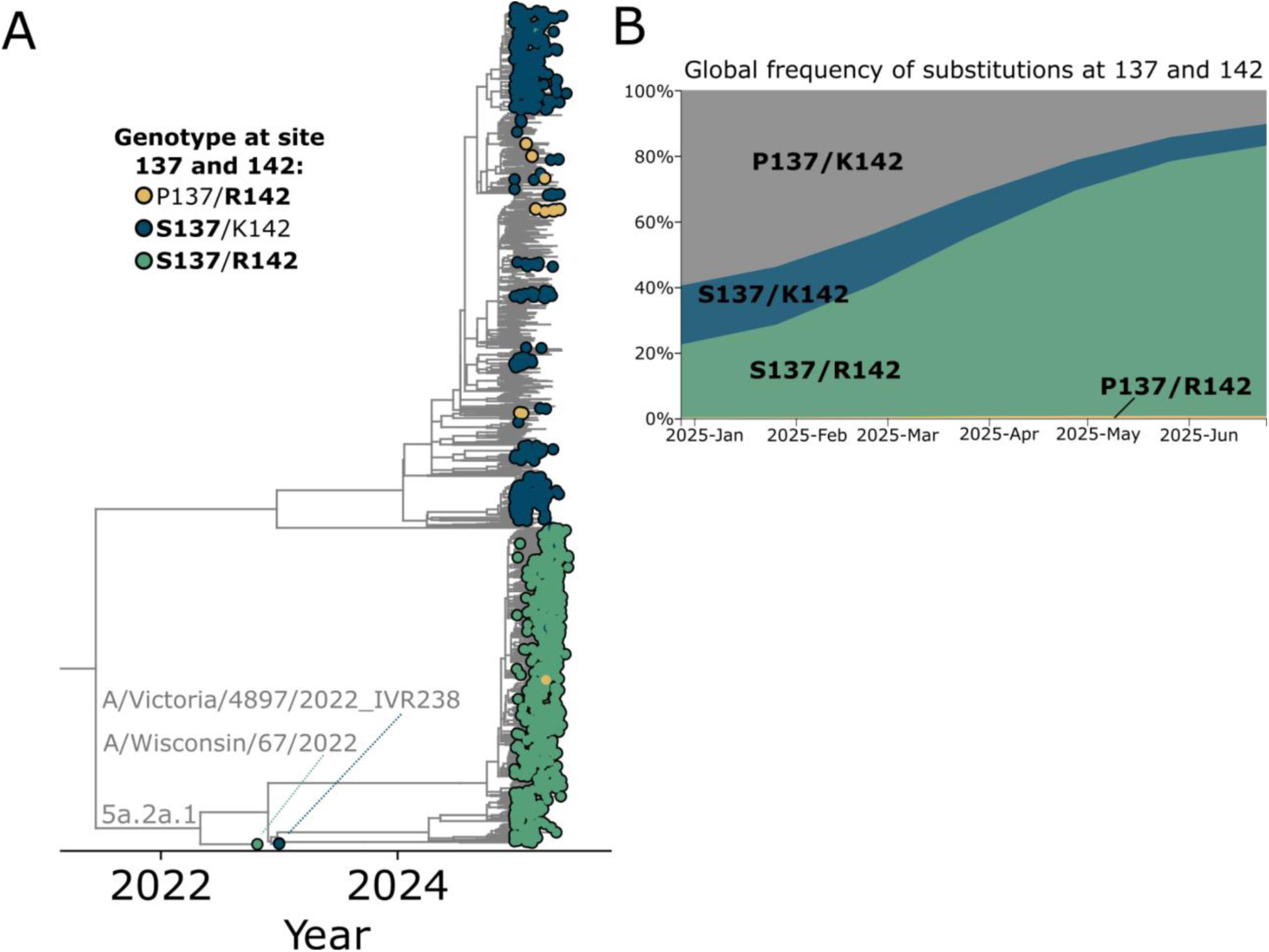
Global frequency of mutations identified to have antigenic effects on pre- and post-vaccination sera first 6 months of 2025. A) A 6-month H1 phylogeny adapted from https://nextstrain.org/seasonal-flu/h1n1pdm/ha/6m@2025-06-26 from Nextstrain^41,42^, showing an increase in strains containing P137S and K142R. Strains containing the K142R mutation only are indicated with yellow circles at the tips, while strains containing P137S only are indicated with navy circles. Strains that contain both P137S and K142R are indicated with green circles. B) The global frequency over time of strains containing mutations P137S and/or K142R.

## References

1. World Health Organization. Seasonal influenza fact sheet. https://www.who.int/news-room/fact-sheets/detail/influenza-(seasonal).

2. Morris, D. H. et al. Predictive Modeling of Influenza Shows the Promise of Applied Evolutionary Biology. Trends Microbiol. 26, 102–118 (2018).

3. Huddleston, J. et al. Integrating genotypes and phenotypes improves long-term forecasts of seasonal influenza A/H3N2 evolution. eLife 9, e60067 (2020).

4. Wong, S.-S. & Webby, R. J. Traditional and New Influenza Vaccines. Clin. Microbiol. Rev. 26, 476–492 (2013).

5. DeMarcus, L., Shoubaki, L. & Federinko, S. Comparing influenza vaccine effectiveness between cell-derived and egg-derived vaccines, 2017–2018 influenza season. Vaccine 37, 4015–4021 (2019).

6. Fowlkes, A. L. et al. Randomised immunogenicity trial comparing 2019-2020 recombinant and egg-based influenza vaccines among frequently vaccinated healthcare personnel in Israel. Int. J. Infect. Dis. 149, 107260 (2024).

7. Rayens, E. et al. Comparative Effectiveness of Cell-Based Versus Egg-Based Influenza Vaccines in Prevention of Influenza Hospitalization During the 2022–2023 Season Among Adults 18–64 Years. Influenza Other Respir. Viruses 18, e70025 (2024).

8. Zost, S. J. et al. Contemporary H3N2 influenza viruses have a glycosylation site that alters binding of antibodies elicited by egg-adapted vaccine strains. Proc. Natl. Acad. Sci. 114, 12578–12583 (2017).

9. Garretson, T. A., Petrie, J. G., Martin, E. T., Monto, A. S. & Hensley, S. E. Identification of human vaccinees that possess antibodies targeting the egg-adapted hemagglutinin receptor binding site of an H1N1 influenza vaccine strain. Vaccine 36, 4095–4101 (2018).

10. Gouma, S. et al. Comparison of Human H3N2 Antibody Responses Elicited by Egg-Based, Cell-Based, and Recombinant Protein–Based Influenza Vaccines During the 2017–2018 Season. Clin. Infect. Dis. Off. Publ. Infect. Dis. Soc. Am. 71, 1447–1453 (2019).

11. Loes, A. N. et al. High-throughput sequencing-based neutralization assay reveals how repeated vaccinations impact titers to recent human H1N1 influenza strains. J. Virol. 98, e00689–24 (2024).

12. Kikawa, C. et al. High-throughput neutralization measurements correlate strongly with evolutionary success of human influenza strains. eLife 14, (2025).

13. Kikawa, C. et al. Near real-time data on the human neutralizing antibody landscape to influenza virus to inform vaccine-strain selection in September 2025. Virus Evol. 11, veaf086 (2025).

14. Recommended composition of influenza virus vaccines for use in the 2023-2024 northern hemisphere influenza season. https://www.who.int/publications/m/item/recommended-composition-of-influenza-virus-vaccines-for-use-in-the-2023-2024-northern-hemisphere-influenza-season (2023).

15. Xu, R. et al. Structural Basis of Preexisting Immunity to the 2009 H1N1 Pandemic Influenza Virus. Science 328, 357–360 (2010).

16. Shu, Y. & McCauley, J. GISAID: Global initiative on sharing all influenza data – from vision to reality. Eurosurveillance 22, 30494 (2017).

17. CDC. Influenza Activity in the United States during the 2023–2024 Season and Composition of the 2024–2025 Influenza Vaccine. Influenza (Flu) https://www.cdc.gov/flu/whats-new/flu-summary-2023-2024.html (2024).

18. Maurel, M. et al. Interim 2023/24 influenza A vaccine effectiveness: VEBIS European primary care and hospital multicentre studies, September 2023 to January 2024. Eurosurveillance 29, 2400089 (2024).

19. Skowronski, D. M., et al. 2023/24 mid-season influenza and Omicron XBB.1.5 vaccine effectiveness estimates from the Canadian Sentinel Practitioner Surveillance Network (SPSN). Eurosurveillance 29, (2024).

20. Chung, J. R. et al. Influenza Vaccine Effectiveness Against Medically Attended Outpatients Illness, United States, 2023–2024 Season. Clin. Infect. Dis. ciae658 (2025) doi:10.1093/cid/ciae658.

21. Wang, W. et al. A mutation in the receptor binding site enhances infectivity of 2009 H1N1 influenza hemagglutinin pseudotypes without changing antigenicity. Virology 407, 374–380 (2010).

22. Yang, L. et al. Mutations associated with egg adaptation of influenza A(H1N1)pdm09 virus in laboratory based surveillance in China, 2009–2016. Biosaf. Health 01, 41–45 (2019).

23. Liu, F. et al. Redirecting antibody responses from egg-adapted epitopes following repeat vaccination with recombinant or cell culture-based versus egg-based influenza vaccines. Nat. Commun. 15, 254 (2024).

24. Liu, F. et al. Influence of Immune Priming and Egg Adaptation in the Vaccine on Antibody Responses to Circulating A(H1N1)pdm09 Viruses After Influenza Vaccination in Adults. J. Infect. Dis. 218, 1571–1581 (2018).

25. FDA. Flulaval Package Insert. https://www.fda.gov/vaccines-blood-biologics/vaccines/flulaval.

26. FDA. Flublok Package Insert. https://www.fda.gov/vaccines-blood-biologics/vaccines/flublok.

27. Tamerius, J. et al. Global Influenza Seasonality: Reconciling Patterns across Temperate and Tropical Regions. Environ. Health Perspect. 119, 439–445 (2011).

28. Yuan, H., Kramer, S. C., Lau, E. H. Y., Cowling, B. J. & Yang, W. Modeling influenza seasonality in the tropics and subtropics. PLoS Comput. Biol. 17, e1009050 (2021).

29. Li, S. H. et al. Childhood immunological imprinting of cross-subtype antibodies targeting the hemagglutinin head domain of influenza viruses. 2025.09.24.25335646 Preprint at 10.1101/2025.09.24.25335646 (2025).

30. Raymond, D. D. et al. Influenza immunization elicits antibodies specific for an egg-adapted vaccine strain. Nat. Med. 22, 1465–1469 (2016).

31. Wilson, J. L. et al. Antigenic alteration of 2017-2018 season influenza B vaccine by egg-culture adaption. Front. Virol. 2, (2022).

32. Katz, J. M. & Webster, R. G. Efficacy of Inactivated Influenza A Virus (H3N2) Vaccines Grown in Mammalian Cells or Embryonated Eggs. J. Infect. Dis. 160, 191–198 (1989).

33. Kilbourne, E. D. et al. Influenza A virus haemagglutinin polymorphism: pleiotropic antigenic variants of A/Shanghai/11/87 (H3N2) virus selected as high yield reassortants. J. Gen. Virol. 74, 1311–1316 (1993).

34. Meyer, W. J. et al. Influence of Host Cell-Mediated Variation on the International Surveillance of Influenza A (H3N2) Viruses. Virology 196, 130–137 (1993).

35. Chen, Z., Zhou, H. & Jin, H. The impact of key amino acid substitutions in the hemagglutinin of influenza A (H3N2) viruses on vaccine production and antibody response. Vaccine 28, 4079–4085 (2010).

36. Bolton, M. J. et al. Antigenic and virological properties of an H3N2 variant that continues to dominate the 2021–22 Northern Hemisphere influenza season. Cell Rep. 39, (2022).

37. Skowronski, D. M. et al. Low 2012–13 Influenza Vaccine Effectiveness Associated with Mutation in the Egg-Adapted H3N2 Vaccine Strain Not Antigenic Drift in Circulating Viruses. PLOS ONE 9, e92153 (2014).

38. Harding, A. T. & Heaton, N. S. Efforts to Improve the Seasonal Influenza Vaccine. Vaccines 6, 19 (2018).

39. Cowling, B. J. et al. Preliminary Findings From the Dynamics of the Immune Responses to Repeat Influenza Vaccination Exposures (DRIVE I) Study: A Randomized Controlled Trial | Clinical Infectious Diseases | Oxford Academic. https://academic.oup.com/cid/article/79/4/901/7718587.

40. Lee, J. M., et al. Deep mutational scanning of hemagglutinin helps predict evolutionary fates of human H3N2 influenza variants. Proc. Natl. Acad. Sci. 115, E8276–E8285 (2018).

41. Hadfield, J. et al. Nextstrain: real-time tracking of pathogen evolution. Bioinformatics 34, 4121–4123 (2018).

42. Hedges, S. B., Dudley, J. & Kumar, S. TimeTree: a public knowledge-base of divergence times among organisms. Bioinformatics 22, 2971–2972 (2006).

43. Bacsik, D. J. et al. Influenza virus transcription and progeny production are poorly correlated in single cells. eLife 12, RP86852 (2023).

44. Welsh, F. C. et al. Age-dependent heterogeneity in the antigenic effects of mutations to influenza hemagglutinin. Cell Host Microbe 32, 1397–1411.e11 (2024).

45. Hoffmann, E., Neumann, G., Kawaoka, Y., Hobom, G. & Webster, R. G. A DNA transfection system for generation of influenza A virus from eight plasmids. Proc. Natl. Acad. Sci. 97, 6108–6113 (2000).

